# Isoform specificity of PKMs during long-term facilitation in *Aplysia* is mediated through stabilization by KIBRA

**DOI:** 10.1101/620633

**Authors:** Larissa Ferguson, Jiangyuan Hu, Diancai Cai, Shanping Chen, Tyler W Dunn, David L Glanzman, Samuel Schacher, Wayne S. Sossin

## Abstract

Persistent activity of protein kinase M (PKM), the truncated form of protein kinase C (PKC), can maintain long-term changes in synaptic strength in many systems including the hermaphrodite marine mollusk, *Aplysia californica.* Moreover, different types of long-term facilitation (LTF) in cultured *Aplysia* sensorimotor synapses rely on the activities of different PKM isoforms in the presynaptic sensory neuron and postsynaptic motor neuron. When the atypical PKM isoform was required, the kidney and brain expressed adaptor protein (KIBRA) was also required. Here, we explore how this isoform specificity is established. We find that PKM overexpression in the motor neuron, but not the sensory neuron, is sufficient to increase synaptic strength and that this activity is not isoform specific. KIBRA is not the rate-limiting step in facilitation since overexpression of KIBRA is neither sufficient to increase synaptic strength, nor to prolong a form of PKM-dependent intermediate synaptic facilitation. However, the isoform specificity of dominant negative (DN)-PKMs to erase LTF is correlated with isoform specific competition for stabilization by KIBRA. We identify a new conserved region of KIBRA. Different splice isoforms in this region stabilize different PKMs based on the isoform-specific sequence of an alpha-helix ‘handle’ in the PKMs. Thus, specific stabilization of distinct PKMs by different isoforms of KIBRA can explain the isoform specificity of PKMs during LTF in *Aplysia*.

**Significance Statement:** Long lasting changes in synaptic plasticity associated with memory formation are maintained by persistent protein kinases. We have previously shown in the Aplysia sensorimotor model that distinct isoforms of persistently active protein kinase Cs (PKMs) maintain distinct forms of long-lasting synaptic changes, even when both forms are expressed in the same motor neuron. Here, we show that, while the effects of overexpression of PKMs in not isoform specific, isoform specificity is defined by a ‘handle’ helix in PKMs that confers stabilization by distinct splice forms in a previously undefined domain of the adaptor protein KIBRA. Thus, we define new regions in both KIBRA and PKMs that define the isoform specificity for maintaining synaptic strength in distinct facilitation paradigms.

## Introduction

The long-term changes in synaptic strength that underlie memory can be maintained through persistently active kinases (Greenberg et al., 1987; Lisman, 1994; Sacktor, 2011; Sossin, 2018). One such kinase is PKM, the truncated form of PKC. PKCs are classified into three families— conventional, novel, and atypical—which differ mainly through the structure of their regulatory domain (Sossin, 2007). Despite not having a regulatory domain, PKMs are similarly classified. While much attention has been given to the role of the atypical PKMζ in memory maintenance (Sacktor, 2011), the fact that the catalytic domains of PKC/PKM families are similar in structure suggests that additional PKM isoforms may perform similar functions. PKMζ knock-out mice, for example, are phenotypically normal due to compensation by the closely related iota isoform (Tsokas et al., 2016). While overexpression of PKMζ is sufficient to both increase synaptic strength (Yao et al., 2008) and to strengthen memory (Shema et al., 2011), the isoform specificity of this ability has not been explored.

In *Aplysia californica*, PKM isoforms from different families are important for the maintenance of distinct forms of synaptic plasticity. The conventional, novel, and atypical PKM families are represented by PKM Apl I, PKM Apl II, and PKM Apl III, respectively (Bougie et al., 2009; Sossin, 2007). A DN approach suggested that, in the postsynaptic neuron, PKM Apl I is required for the maintenance of non-associative LTF, while PKM Apl II and PKM Apl III are required for the maintenance of associative LTF (Hu et al., 2017a; Hu et al., 2017b). Because associative and non-associative LTF can co-exist in one postsynaptic cell (Hu et al., 2017b), the cell must have some way of targeting different PKM isoforms to different synapses that underlie these forms of LTF. We propose stabilization by KIBRA as a mechanism through which PKM isoform specificity is achieved.

KIBRA has been implicated in human episodic memory in a number of genome-wide association studies (Milnik et al., 2012; Papassotiropoulos et al., 2006), and co-localizes with and stabilizes the mammalian PKMζ (Vogt-Eisele et al., 2014; Yoshihama et al., 2009). KIBRA regulates AMPA receptor trafficking and is required for late long-term potentiation (LTP) and some forms of memory in rodents (Heitz et al., 2016; Makuch et al., 2011; Vogt-Eisele et al., 2014). In *Aplysia*, KIBRA stabilizes overexpressed PKM Apl II and PKM Apl III, but not PKM Apl I (Hu et al., 2017b). KIBRA-AAA in which the three residues found to be essential for PKMζ binding were mutated (Vogt-Eisele et al., 2014) did not stabilize PKM Apl III (Hastings et al., 2018; Hu et al., 2017b) and erased the forms of LTF blocked by DN PKM Apl III (Hu et al., 2017b), but did stabilize PKM Apl I and did not erase the forms of LTF blocked by DN PKM Apl I. However, whether DN PKMs act by competition for KIBRA is unclear and indeed, in vertebrates, KIBRA is unable to bind a catalytically inactive form of PKMζ (Vogt-Eisele et al., 2014) and the DN PKMs used in *Aplysia* are catalytically inactive (Bougie et al., 2012).

Here, we explore the sufficiency of PKMs and KIBRA to increase synaptic strength and the role of KIBRA in determining isoform specificity of PKMs. Overexpression of PKMs in the motor neuron, but not the sensory neuron, increased synaptic strength. Unlike PKMs, KIBRA is not sufficient to increase synaptic strength when overexpressed in the motor neuron. However, DN PKMs competed for isoform-specific stabilization by KIBRA and alternatively spliced forms of KIBRA stabilized specific isoforms of PKM based on the sequence in their alpha-helix ‘handle’ domain. Thus, isoform specificity of PKMs is explained by differential stabilization by KIBRA isoforms.

## Materials and Methods

### Aplysia cell culture

*Aplysia californica* from Miami *Aplysia* Resource Facility (RSMAS) or from Alacrity marine biological services (Redondo Beach CA, USA) were anesthetized via injection of 50-60 mL of 400 mM isotonic MgCl_2_ and abdominal and/or pleuropedal ganglia were removed. Ganglia were digested at 19°C in L15 media containing 10 mg/ml Dispase II (Roche Diagnostics) for 18-19 h or for experiments from animals from Alacrity at 35°C for 2 h. L15 medium (Sigma-Aldrich) was supplemented with 0.2 M NaCl, 26 mM MgSO_4_, 35 mM dextrose, 27 mM MgCl_2_, 4.7 mM KCl, 2 mM NaHCO_3_, 9.7 mM CaCl_2_, and 15 mM HEPES, with pH 7.4. Glass bottom culture dishes were coated with 0.05% poly-L-lysine for 1-2 h and washed with ddH_2_O prior to use. Sensory neurons and LFS motor neurons were isolated from pleural and abdominal ganglia respectively that were dissected from adult *Aplysia* (60–100g) and L7 motor neurons were isolated from the abdominal ganglia of juvenile animals (2 g). Neurons were cultured in 50% hemolymph/50% L15 media supplemented with L-glutamine. For electrophysiology experiments, motor neurons were removed from the abdominal ganglia and allowed to adhere to the culture dish for 1-24 h before pairing with a sensory neuron from pleural ganglia as previously described (Zhao et al., 2006). Note that each coculture comprised a single presynaptic sensory neuron paired with a single postsynaptic motor neuron (either an LFS or an L7 neuron). Cells were incubated for 48 h at 19°C (SN-LFS) to allow time for them to adhere to the dish prior to injection or for 96 h at 19°C (SN-L7) to allow time for the formation of stable synapses (Hu and Schacher, 2015; Hu et al., 2017a, 2017b). All plating, injections, and electrophysiology experiments were performed at 19°C, except for experiments using animals from Alacrity which were performed at room temperature.

### Plasmid constructs and microinjection

All constructs were made in the pNEX3 vector (Kaang, 1996) and for all vector experiments pNEX3 plasmid was used. All PKM constructs were made as fusion proteins with monomeric red fluorescent protein (mRFP), while PKC Apl I was a fusion protein with enhanced green fluorescent protein (eGFP). The mRFP or eGFP has been removed from the construct names for clarity. Dominant negative constructs (DN PKM Apl I, DN PKM Apl II, and -DN PKM Apl III) and PKCs/PKMS used for overexpression and stabilization studies (PKC Apl I, PKM Apl I, PKM Apl II and PKM Apl III) were previously described (Bougie et al., 2012; Farah et al., 2017; Hu et al., 2017a). The new DN PKM Apl III K-R was generated by cutting out this region from the plasmid encoding PKC Apl III K-R (Bougie et al, 2012) with AarI and SalI and inserting into the plasmid encoding PKM Apl III plasmid at the same sites. For the chimeras, GBLOCKS (Integrated DNA technology, IO, USA) were purchased with the PKM Apl III sequences [carboxy-terminus (CT) or handle] replaced by PKM Apl I sequences. These were then cut out with either BsmBI and Kpn I(CT) or Sal I and Kpn I (handle) and inserted into the plasmid encoding PKM Apl III at the same sites. The chimeras were sequenced for confirmation. Plasmids encdoding KIBRA and KIBRA-AAA were previously described (Hu et al., 2017b) and are not fusion proteins with a fluorescent protein. The KIBRA splice form, KIBRA SPL, was generated similarly to KIBRA but from a separate PCR clone that fortuitously encoded the splice form (Hu et al., 2017b). For stabilization experiments, molar equivalent levels of plasmids encoding KIBRA/KIBRA SPL/KIBRA-AAA and the PKMs/DN PKMs were used. A solution containing the desired constructs (max 0.4 ug/ul of DNA in ultrapure water) with 0.2% fast green were microinjected into the nuclei of neurons using back-filled glass micropipettes. A short pressure pulse was delivered to release plasmid solution into nucleus. KIBRA and enhanced green fluorescent protein (eGFP) were injected into the LFS neuron of sensorimotor cocultures for ITF experiments. KIBRA was co-expressed with an eGFP construct so that expression could be confirmed by detecting eGFP fluorescence. Stabilization experiments were performed by injecting constructs into isolated sensory neurons. For PKM overexpression assays in sensorimotor cocultures, the constructs were injected after monitoring EPSP amplitudes on day 0. Injection into L7 motor neurons was performed as previously described (Hu et al., 2017b). Cultures were incubated at 19 ºC for 24 h to allow sufficient time for expression of injected plasmids prior to imaging.

### Electrophysiology

For sensorimotor cocultures containing L7 motor neuron, the amplitude of the excitatory postsynaptic potential (EPSP) was recorded as previously described (Hu et al., 2017b). The assignment of any culture to control and experimental groups at the start of each experiment on day 4 (defined as day 0 in Results, Figure 1 and Figure 3) was to ensure that there were no significant differences in the synaptic strength between the groups before treatments. Sensorimotor cocultures containing LFS motor neurons were incubated in culture media at room temperature for 24 h following microinjection to allow sufficient time for expression of injected plasmids before electrophysiological recordings. Prior to recording, culture media was replaced with a recording saline [NaCl (460 mM); MgCl_2_ (55 mM); CaCl_2_ (10 mM); KCl (10 mM); D-Glucose (10 mM); HEPES (10 mM); pH 7.6]. Membrane potentials were recorded and controlled in current clamp mode with sharp intracellular electrodes attached to an Axoclamp 2B amplifier (Molecular Devices, Palo Alto, California). Microelectrodes (15-30 MΩ) were backfilled with 2 M potassium acetate and bridge-balanced before and after membrane penetration. The postsynaptic cell was impaled first, so that if entry into the presynaptic cell resulted in generation of an action potential, the resultant postsynaptic potential (PSP) would be recorded. Injection of hyperpolarizing current was used to keep both presynaptic and postsynaptic neurons at −80 mV during recording. Postsynaptic input resistance was measured with 500 msec, −0.5 nA pulses. For ITF experiments, electrodes were removed from cells after initial recording and 5HT (10 µM) was added for 10 min before being washed out with 25 ml recording saline solution. Cultures were left at 19ºC for 2 hours, at which time the second recording was performed. Initial PSP rise-rate was measured as previously described (Dunn and Sossin, 2013). For PKM overexpression experiments (Figure 1), EPSP amplitudes were measured prior to microinjection (day 0 = 96 h after cell plating to insure stable synapses), and again on days 1, 3, and 5 as described (Bougie et al., 2012; Hu et al., 2017a). The kinetics of homosynaptic depression (HSD) were monitored on day 1 by homosynaptic stimulation— stimulating the sensory neuron with low frequency stimulation (one action potential every 20 seconds; 6 stimuli in total)(Hu et al., 2017b). The motor neuron was maintained at −80 mV during the stimulation to accurately record the amplitude of each EPSP. The initial EPSP amplitude evoked by the first stimulus was normalized as 100%.

**Figure 1.**
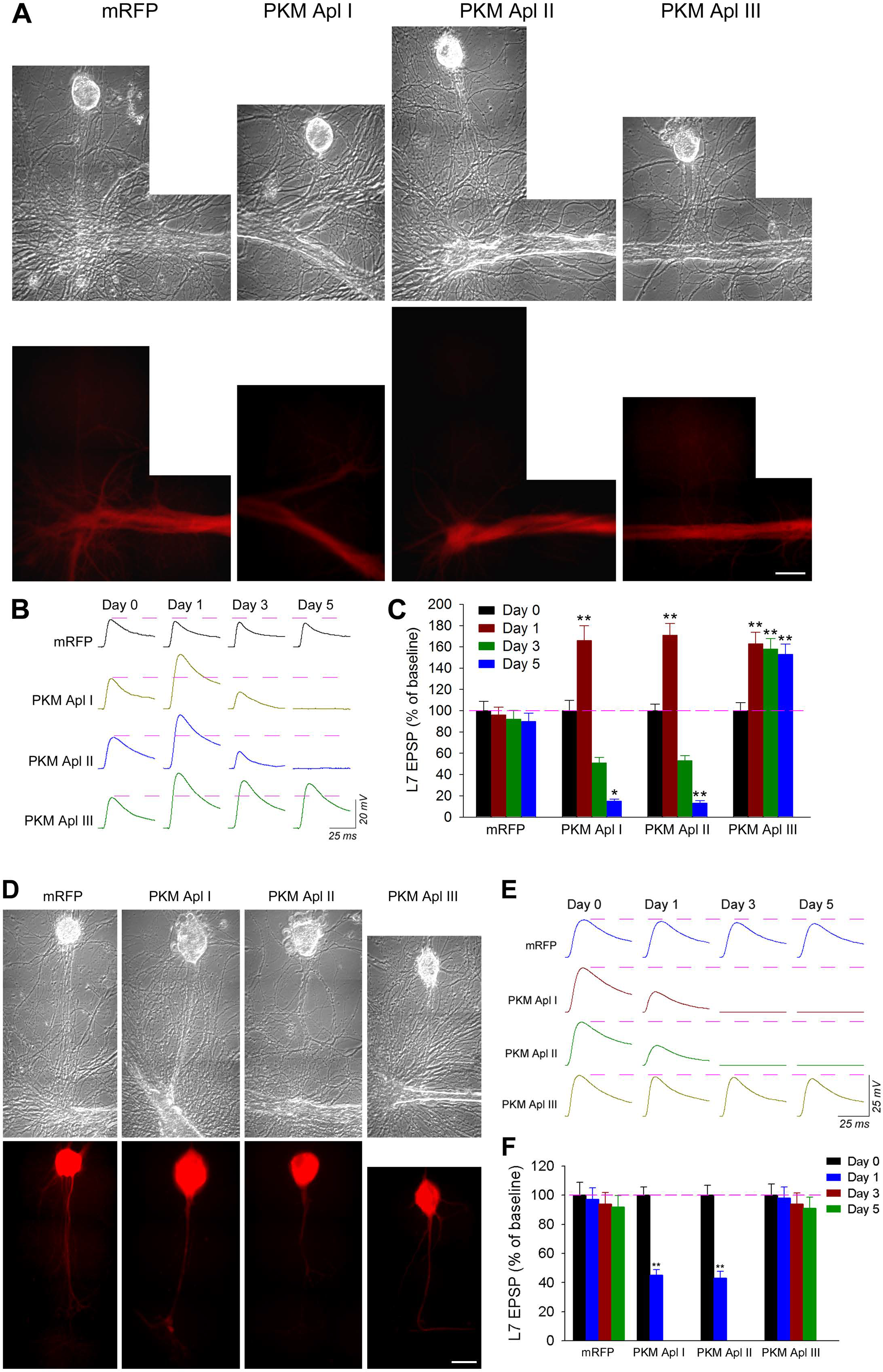
Pre- and post-synaptic overexpression of PKM isoforms produce different changes in synaptic strength. **A**, Phase contrast and epifluorescent views of co-cultures 24 hours after the indicated construct was injected into each postsynaptic neuron L7. The constructs were injected after monitoring EPSP amplitudes on day 0 (96 h after plating sensory neuron and L7). The scale bar equals 25µm. **B**, Sample EPSP traces for each treatment. The dashed lines represent the synaptic baseline. EPSPs recorded from cultures before (day 0) and at various times after injection of the indicated construct. Overexpression of PKM Apl III in L7 produced a significant increase in synaptic baseline compared to control (overexpressed mRFP) and persisted to day 5. Overexpression of PKM Apl I or PKM Apl II also produced a significant increase in EPSP amplitude on day 1, but EPSP amplitude declined after day 3. **C**, Summary of the change in EPSP amplitudes (% of baseline). A two-way ANOVA indicated a significant effect of group X repeated measures (*F*_9, 69_ = 104.831, *p*< 0.0001). Pairwise comparisons (Bonferroni) indicated that overexpression of PKMApl III in L7 (n = 14) significantly increased EPSP amplitudes on day 1, 3 and 5 compared to mRFP (n = 8; \*\**p*< 0.01). Overexpression of PKMApl I (n = 11) or PKM Apl II (n= 9) significantly increased EPSP amplitudes on day 1 compared to mRFP (\*\**p*< 0.01), but EPSP amplitudes declined after day 3 and significantly decreased on day 5 (\**p*< 0.05; \*\**p*< 0.01) compared to mRFP. On day 5, only 3 of 11 cultures with overexpression of PKM Apl I had small EPSPs, the resting membrane potentials of L7 in the other 8 cultures without EPSPs were close to zero, while 2 of 9 cultures with overexpression of PKM Apl II had small EPSPs and the other 7 cultures that lacked EPSPs had resting potentials close to zero. Error bars represent SEM. **D**, Phase contrast and epifluorescent views of co-cultures 24 hours after the indicated construct was injected into each SN. The constructs were injected after monitoring EPSP amplitudes on day 0 (96 h after plating). The scale bar equals 25 µm. **E**, Sample EPSP traces for each treatment. The dashed lines represent the synaptic baseline. EPSPs recorded from cultures before (day 0) and at various times after injection of the indicated construct. Overexpression of PKM Apl III had no significant impact on synaptic baseline compared to control (mRFP overexpression). In contrast, overexpression of PKM Apl I or PKM Apl II produced a significant decline in EPSP amplitude on day 1. Viability of sensory neurons (extensive degeneration of cell body and neurites) was compromised by day 3 resulting in an absence of EPSPs. **F**, Summary of the change in EPSP amplitudes (% of baseline). Overexpression of PKM Apl III in SN (n = 11) evoked no change in synaptic baseline compared to control (n = 7), while overexpression of PKM Apl I (n = 11) or PKM Apl II (n = 10) produced a significant decline in EPSP and subsequent death of the sensory neuron. A two-way ANOVA indicated a significant effect of group X repeated measures (*F*_9, 105_ = 100.507, *p*< 0.0001). Pairwise comparisons (Bonferroni) indicated that overexpression of PKM Apl III evoked no significant change on day 1, 3 and 5 compared to Control (all *p*> 0.9). Overexpression of PKM Apl I or PKM Apl II produced a significant decline in EPSP amplitudes on day 1 (all \*\**p*< 0.01). Error bars represent SEM.

### Immunocytochemistry

Cultured cells were fixed with 4% paraformaldehyde + 30% sucrose in PBS for 30 min and washed with PBS. Fixed cells were permeabilized (0.1% Triton X-100 with 30% sucrose in PBS for 10 min) and free aldehydes were quenched (50 mM ammonium chloride for 10 min, followed by four PBS washes). Cells were incubated with 10% normal goat serum (Sigma) + 0.5% Triton X-100 in PBS for 30 min to block nonspecific antibody binding. Cells were then treated with PKC Apl III antibody in blocking solution (1:5000) for 1 h, followed by five PBS washes (10 min each). Cells were then incubated with Alexa Fluor 647-conjugated donkey anti-rabbit antibody in blocking solution (1:500, Invitrogen) for 1 h in darkness and washed with PBS as described above.

### Imaging and image analysis

Images were captured by an LSM 710 (Zeiss) laser confocal scanning microscope equipped with an Axiovert 100 inverted microscope (Zeiss) and a 40×, NA 1.4 objective. The eGFP, mRFP, and/or A647 images were acquired sequentially. Images for the expression of PKM constructs in sensorimotor cocultures were viewed with a Nikon Diaphot microscope attached to a silicon-intensified target (SIT) (Dage 68; Dage-MTI) video camera. Fluorescence intensity (arbitrary units) was measured by the Microcomputer-Controlled Imaging Device (MCID) software package (Imaging Research). For analysis of DN PKM and PKM stabilization, single processes for each neuron were outlined using NIH Image J. Background fluorescence was subtracted from the fluorescence values measured. The red/green or cyan/green ratio for all neurons was normalized to the average ratio seen in vector-injected neurons from the same experiment. For most stabilization assays, a one-way ANOVA of the normalized ratio was performed with Tukey post hoc tests. When only two groups were present, a Student’s T-test was used. If multiple comparisons were made for two groups (as in the experiments with chimeric PKMs), a Bonferonni correction for multiple tests was used. All quantification of stabilization was done blindly.

#### Evolution analysis

Orthologues of KIBRA were determined using the reverse BLAST method as described below. Organisms with established genomes on NCBI were searched using BLAST, to identify proteins with homology to KIBRA. To distinguish between true orthologues and proteins with similar WW and C2 domains, these proteins were then used as a query in a BLAST search and if they are much more homologous to other proteins with WW and C2 domains than to KIBRA, they were rejected as orthologues. Based on this, no KIBRA orthologues exist in sponge (*Amphimedon queenslandia*), Trichoplax (*Trichoplax adhaerens*) and choanoflagellate (*Monosiga bervicolis*; *Salpingoeca rosetta*). KIBRA orthologues were found in other Cnidaria (Coral, *Styklophora postillata* and *Acropora digitefara*; jellyfish, *Hydra vulgaris)*, but the strongest homology was to Anemone, *Nemostella vectensis*, and this Cnidarian was used in the comparison studies.

### Behavioral training

Adult *Aplysia* (80-120 g) were given 1 d of long-term sensitization training. The training consisted of 5 bouts of tail shocks delivered via implanted platinum wires. Each bout comprised three trains (train duration = 1 s) of shocks (10-ms pulse duration, 40 Hz, 120 V) spaced 2 s apart; the bouts of shocks were separated by 20-min intervals. Prior to the shocks, each animal was given three pretests, spaced 10-min apart, during which a brief, relatively weak tactile stimulus was applied to its siphon; the duration of the resulting siphon-withdrawal response (SWR) providing an index of sensitization. Forty-eight hours after the sensitization training, each snail was given a single test stimulus, and its SWR measured. Another group of snails were treated identically to the trained group, except that the tail shocks on day 1 were withheld. Animals that received the shock training exhibited a significantly prolonged SWR compared to animals that received only the test stimuli (Chen et al., 2014). Immediately after the 48-h test, the animals were anesthetized by immersion in cold (4°C) seawater, and their abdominal ganglia were dissected out and prepared for QRT-PCR.

### QRT-PCR

Total RNA was extracted from abdominal ganglia of trained and control (test alone) *Aplysia* using Trizol reagent following the manufacturer’s instructions. cDNA was synthesized from 2 μg total RNA using qScript™ cDNA SuperMix, (QuantaBio) to a final concentration of 100 ng/μl. KIBRA primer pair sequences were used: forward “TGG AGA AAC TCC TGC AAG GCC A”; rev “ATT GCT GGC GTC TGC TAA CTC”. Quantitative RT-PCR was performed using the ABI PRISM 7900 sequencing detection system (Applied Biosystems, Foster City, CA, USA) with SYBR green master mix under the following conditions: initial denaturation at 95°C for 10 min, followed by 40 cycles of 95°C for 15 s and 65°C for 30 s, 72° 30 s. 5 μl SYBR, 3 μl H2O, 1 μl cDNA per reaction, each sample was run in triplicate. The data were quantified by the 2(-Delta Delta C(T)) method (Livak and Schmittgen, 2001). H4 was used as an internal control.

## Results

### Postsynaptic, but not presynaptic, overexpression of PKM is sufficient to produce a long-lasting increase in synaptic strength

In vertebrates, overexpression of PKMζ is sufficient to increase synaptic strength when expressed in the postsynaptic neuron (Yao et al., 2008). However, there have been no studies examining the isoform specificity of this effect or whether PKMs are sufficient to increase synaptic strength when overexpressed in the presynaptic neuron. Overexpression of any of the three *Aplysia* PKM isoforms in the postsynaptic motor neuron of *Aplysia* sensorimotor co-cultures is sufficient to increase synaptic strength 24 h following injection compared to overexpression of mRFP alone (Fig. 1A-C). Continued expression of PKM Apl I or PKM Apl II is toxic while overexpression of PKM Apl III can maintain increased synaptic strength for a prolonged period (Fig. 1A-C). In contrast, none of the PKMs increased synaptic strength 24 h following injection when expressed in the sensory neuron (Fig.1D-F). Similar to the motor neuron, overexpression of PKM Apl I and PKM Apl II in the sensory neuron was toxic, but PKM Apl III was not (Fig. 1D-F). These results were obtained using the postsynaptic neuron L7 (gill withdrawal motor neuron), but similar results were seen for PKM Apl III in the presynaptic and postsynaptic neuron when LFS neurons (siphon withdrawal motor neurons) were used as the postsynaptic neurons (Fig. 2).

**Figure 2.**
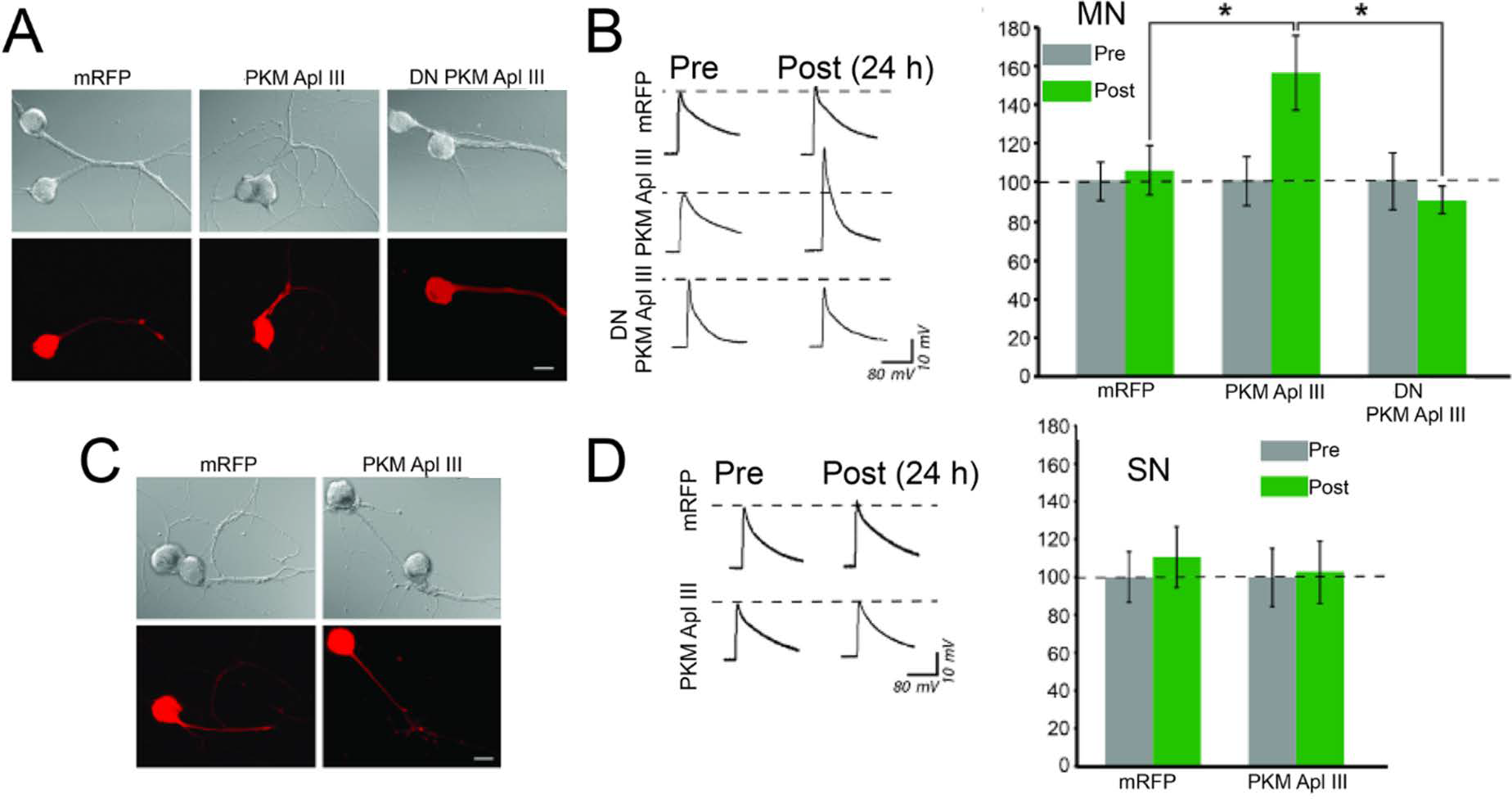
Overexpression of PKM Apl III in the LFS motor neurons, not in the sensory neurons, produces a long-term increase in synaptic strength. **A)** and **C)** Phase contrast and epifluorescent images of the same view areas of LFS motor neurons (A) or sensory neurons (C) expressing mRFP or PKM Apl III 24 hours after injection of constructs. The constructs were injected into the sensory or the motor neurons on day 4 of cell culturing. The bar is 20 µm. **B)** Overexpression of PKM Apl III, but not DN PKM Apl III in LFS motor neurons produces an increase in synaptic strength. Sample traces show EPSPs evoked in the LFS motor neuron before (pre) and 24 h after (post) control and injections of constructs in the motor neuron. The accompanying histogram is a summary of the change in the EPSP amplitudes. The mean normalized EPSP in the mRFP group (n = 28, 12 independent experiments), PKM Apl III group (n = 22, 12 independent experiments), and DN PKM Apl III group (n = 12, 6 independent experiments), was 106% ± 13%, 154% ± 19%, and 90% ± 7%, respectively. A one-way ANOVA indicated that the overall group differences were significant (F[2,59] = 4.4, p = 0.02). SNK posthoc tests showed that the mean normalized EPSP in the PKM Apl III group was significantly higher than that in the mRFP and DN -PKM Apl III groups, as indicated by the asterisks (P < 0.05 for both comparisons). The difference between the mRFP and DN PKM Apl III groups was not significant. **D)** Overexpression of PKM Apl III in sensory neurons does not affect synaptic strength. Sample traces show EPSPs evoked in the LFS motor neuron before (pre) and 24h after (post) control and injections of constructs in the sensory neuron. The accompanying histogram is a summary of the change in the EPSP amplitudes. There was no effect of presynaptic overexpression of PKM Apl III. The mean normalized EPSP in the PKM Apl III group (n = 9, 6 independent experiments) and mRFP group (n = 9, 5 independent experiments) was 103% ± 17% and 111% ± 16%, respectively.

Associative, but not non-associative LTF, attenuates HSD (Hu and Schacher 2015). DN PKM Apl III erases associative LTF, but not non-associative LTF (Hu et al, 2017b). However, overexpression of PKM Apl III in the postsynaptic neuron, while sufficient to increase synaptic strength, is not sufficient to attenuate HSD (Fig. 3A-C). Since PKM expression in the sensory neuron is toxic (Fig. 1D-F), we determined the effect of overexpression of PKC Apl I in the sensory neuron. While this was not sufficient to increase synaptic strength or to attenuate HSD on its own, in conjunction with PKM Apl III expression in the motor neuron, attenuation of HSD was observed (Fig.3A-C). Thus, the full expression of associative LTF requires kinase activity in both the pre- and post-synaptic neuron.

**Figure 3.**
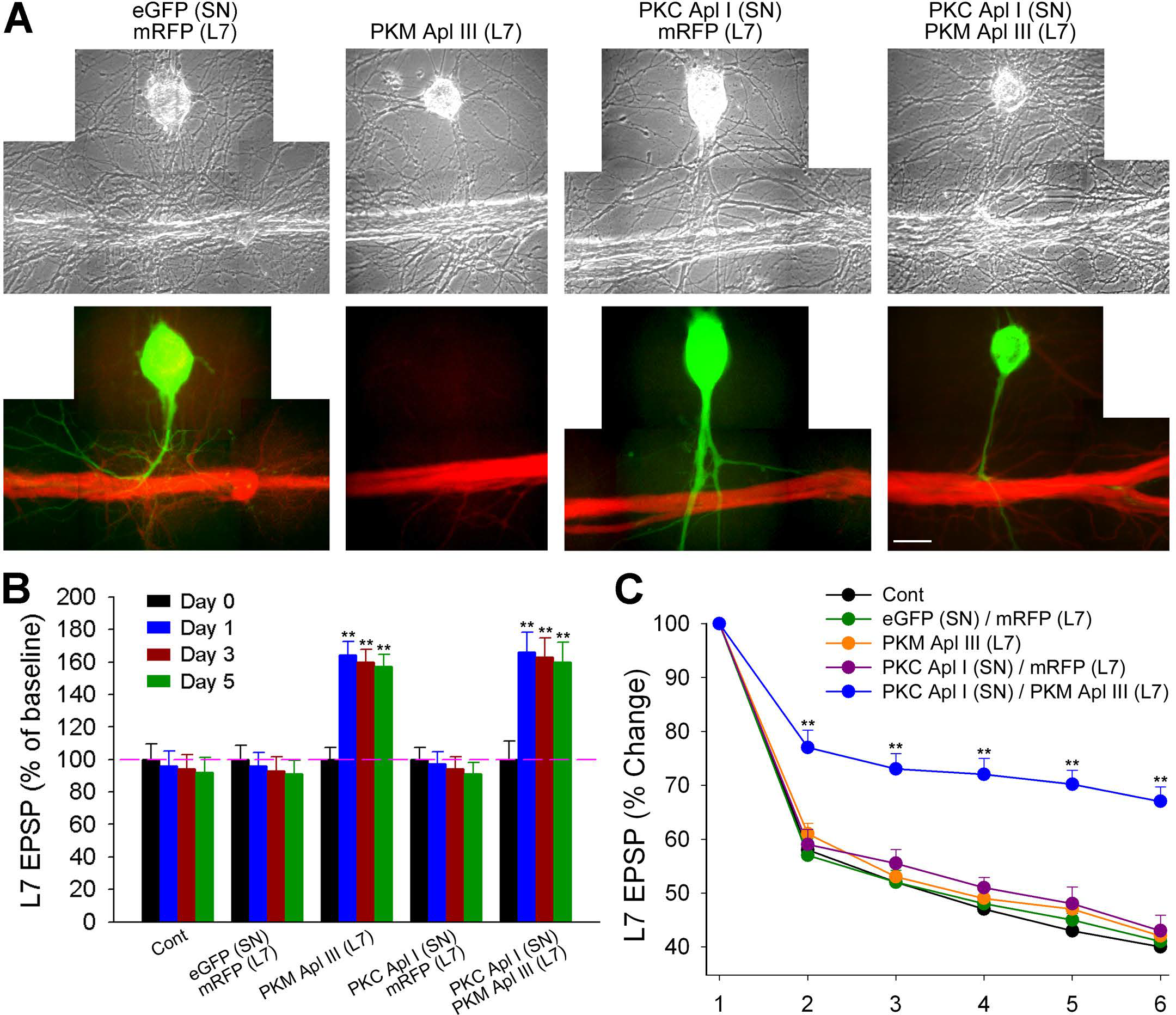
Full expression of associative LTF requires kinase activity in both pre- and post-synaptic neurons. **A)** Phase contrast and epifluorescent views of co-cultures 24 hours after the indicated construct injection in SN and L7. The scale bar equals 25 µm. **B**, Summary of the change in EPSP amplitudes (% of baseline). Overexpression of PKM Apl III alone in L7 (n = 12) or co-overexpression of PKC Apl I in SN and PKM Apl III in L7 (n = 10) produced a persistent increase in the synaptic strength. The other over-expressions failed to increase synaptic strength and were not significantly different than control. A two-way ANOVA indicated a significant effect of group X repeated measures (*F*_12, 108_ = 65.656, *p*< 0.0001). Pairwise comparisons (Bonferroni) indicated that overexpression of either PKM Apl III alone in L7 or both PKC Apl I in SN and PKM Apl III in L7 evoked significant facilitation on day 1, 3 and 5 compared to control (n = 6), co-overexpression of eGFP in SN and mRFP in L7 (n = 7), or co-overexpression of PKC Apl I in SN and mRFP in L7 (n = 6) (all \*\**p*< 0.01). Error bars represent SEM. **C**) The kinetics of HSD after co-overexpression of PKC Apl I in SN and PKM Apl III in L7 was attenuated. Compared to control, only co-overexpression of PKC Apl I in SN and PKM Apl III in L7 evoked an attenuated kinetics of HSD that is observed at sensorimotor synapses expressing persistent associative LTF, while other over expressions failed to alter HSD kinetics. A two-way ANOVA indicated a significant effect of group × repeated measures on the kinetics of HSD (*F*_20, 180_ = 15.025; *p* < 0.0001). Pairwise comparisons (Bonferroni) indicated that the kinetics of HSD only with co-overexpression of PKC Apl I in SN and PKM Apl III in L7 was significantly different than that for control (\*\**p* < 0.01). Error bars represent SEM.

### KIBRA overexpression is not sufficient to prolong ITF

Forms of facilitation that last longer than short-term facilitation but do not require transcription are termed intermediate facilitation (ITF) in *Aplysia*. There are different forms of ITF based on the stimulation, but massed ITF (m-ITF) is of particular interest as it requires cleavage of PKC Apl III to PKM Apl III since: (i) it is blocked by DN PKM Apl III, (ii) it is blocked by a DN form of the protease that cleaves PKCs to PKMs, DN classical calpain, and (iii) stimulation that induces m-ITF causes cleavage of a PKC Apl III FRET construct in the motor neuron (Bougie et al., 2012; Farah et al., 2017). Despite the requirement for a PKM, this form of facilitation is transient (McCamphill et al., 2015; Sutton et al., 2002). One hypothesis is that PKMs are made but not stabilized in the absence of KIBRA and thus transcriptional upregulation of KIBRA could be important for the prolongation of PKM activity and increases in synaptic strength. Indeed, KIBRA stabilizes PKM Apl III (Hu et al, 2017) and here we show that long-term behavioural sensitization in *Aplysia* resulted in an upregulation of KIBRA mRNA as measured by quantitative RT-PCR (Fig. 4A). To determine if increased levels of KIBRA are sufficient to maintain memories by stabilizing PKMs, we induced m-ITF in neurons expressing KIBRA. KIBRA did not, however, increase m-ITF at 2h after induction, a time in which m-ITF has decayed to baseline (McCamphill et al., 2015; Sutton et al., 2002) (Fig. 4B). This was not due to an occlusion by KIBRA as there was no increase in basal synaptic strength in the KIBRA expressing neurons (control 35.99 ± 12.8 mV; KIBRA 30.31 ± 7.4 mV; unpaired t-test p = 0.6919).

**Figure 4.**
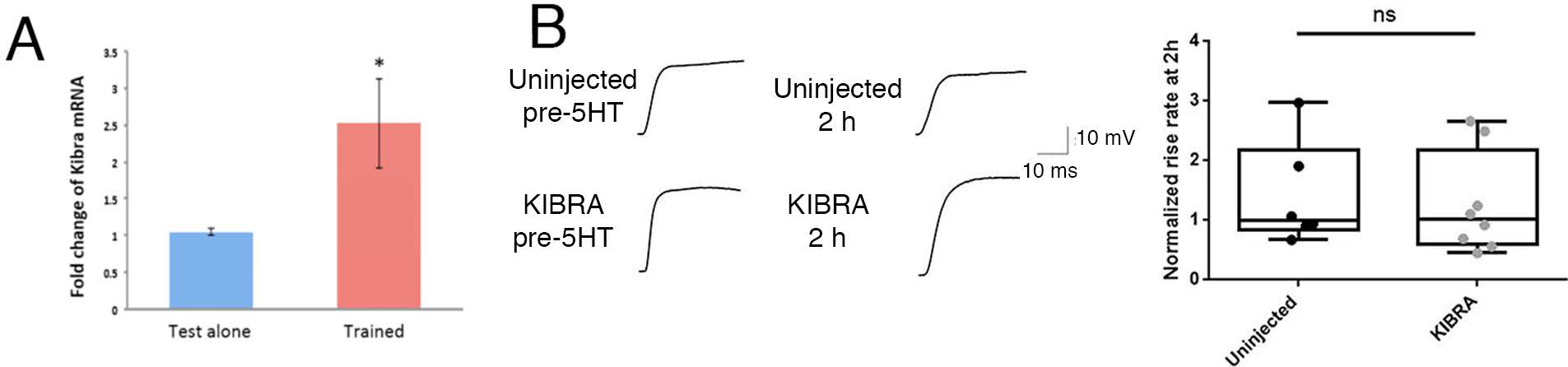
KIBRA overexpression is not sufficient to prolong PKM Apl III-dependent ITF. **A**, QRT-PCR results comparing levels of KIBRA mRNA from cDNA generated from control abdominal ganglia (8 animals) or abdominal ganglia from sensitized animals (7 animals). Results are SEM. *, p<0.05 One-tailed t-test. **B**, Representative traces before and 2 hours after 10 min 10µM 5HT treatment in sensorimotorcocultures overexpressing KIBRA (with eGFP to confirm expression) in the postsynaptic neuron or uninjected controls. Scale bar is 10mV/10ms. **B**, Box and whisker graph with overlaid individual data points (n=6 for uninjected controls and n=8 for KIBRA overexpressing synapses) showing the change in rise-rate 2 hours post-5HT treatment as a percentage of the initial rise rate of the synapses measured. Rise-rate was not significantly different between groups (one-way unpaired t-test [ns, p = 0.7613]). The change in PSP at 2 h was similar between groups via unpaired t-test (ns, p = 0.4380). Initial postsynaptic input resistance and the change in postsynaptic input resistance was similar between groups (180.0 ± 27.4 MΩ uninjected and 177.9 ± 35.6 MΩ eGFP + KIBRA, P=0.9661 that changed to 92.4 ± 6.0 % and 78.6 ± 6.2 % respectively at 2h, p = 0.1448 compared with unpaired t-tests).

### Competition for stabilization by KIBRA accounts for isoform-specific effects of dominant negative PKMs

Since overexpression of any PKM in the motor neuron is sufficient to increase synaptic strength (transiently by PKM Apl I and PKM Apl II and persistently by PKM Apl III), how can distinct PKMs be required in the motor neuron for different forms of LTF when they are both expressed in the same post-synaptic neuron (Hu et al, 2017b)? One possibility is isoform-specific stabilization at synapses by the adaptor protein KIBRA. *Aplysia* KIBRA stabilizes PKM Apl II and PKM Apl III, but not PKM Apl I (Hu et al, 2017b). Similarly, DN PKM Apl II and DN PKM Apl III erase associative LTF when expressed in the motor neuron, as does a form of KIBRA (KIBRA-AAA) that does not stabilize PKM Apl III (Hu et al, 2017b). Thus, DN PKMs may work by competing with endogenously produced PKMs for KIBRA-mediated stabilization. One issue with this proposal is the report that vertebrate KIBRA does not bind to catalytically inactive PKMs (Vogt-Eisele et al., 2014) and the DN PKMs used in this study are catalytically inactive. However, in *Aplysia* neurons, KIBRA stabilizes the DN PKMs (Fig. 5A-B). The pattern of stabilization between KIBRA and KIBRA-AAA was similar to that seen with catalytically active PKMs (Hu et al., 2017b)—KIBRA stabilizes DN PKM Apl III and DN PKM Apl II, whereas KIBRA-AAA stabilizes DN PKM Apl I. These DN PKMs were generated by a aspartate to alanine (D-A) mutationat the catalytic aspartic acid (Cameron et al., 2009) which receives priming phosphorylation (Bougie et al., 2012; Cameron et al., 2009); by contrast, the kinase dead mutant used in Vogt-Eisele et al, 2014—produced through the mutation of a lysine in the ATP binding pocket—results in a PKC that lacks priming phosphorylation (Cameron et al., 2009). However, PKM Apl III with a similar lysine mutation that also does not receive priming phosphorylation (Bougie et al, 2012) was also stabilized by KIBRA (Fig 5C-D). Thus, in *Aplysia*, PKMs do not need to be catalytically active or have priming phosphorylation in order for KIBRA-mediated stabilization to occur.

**Figure 5.**
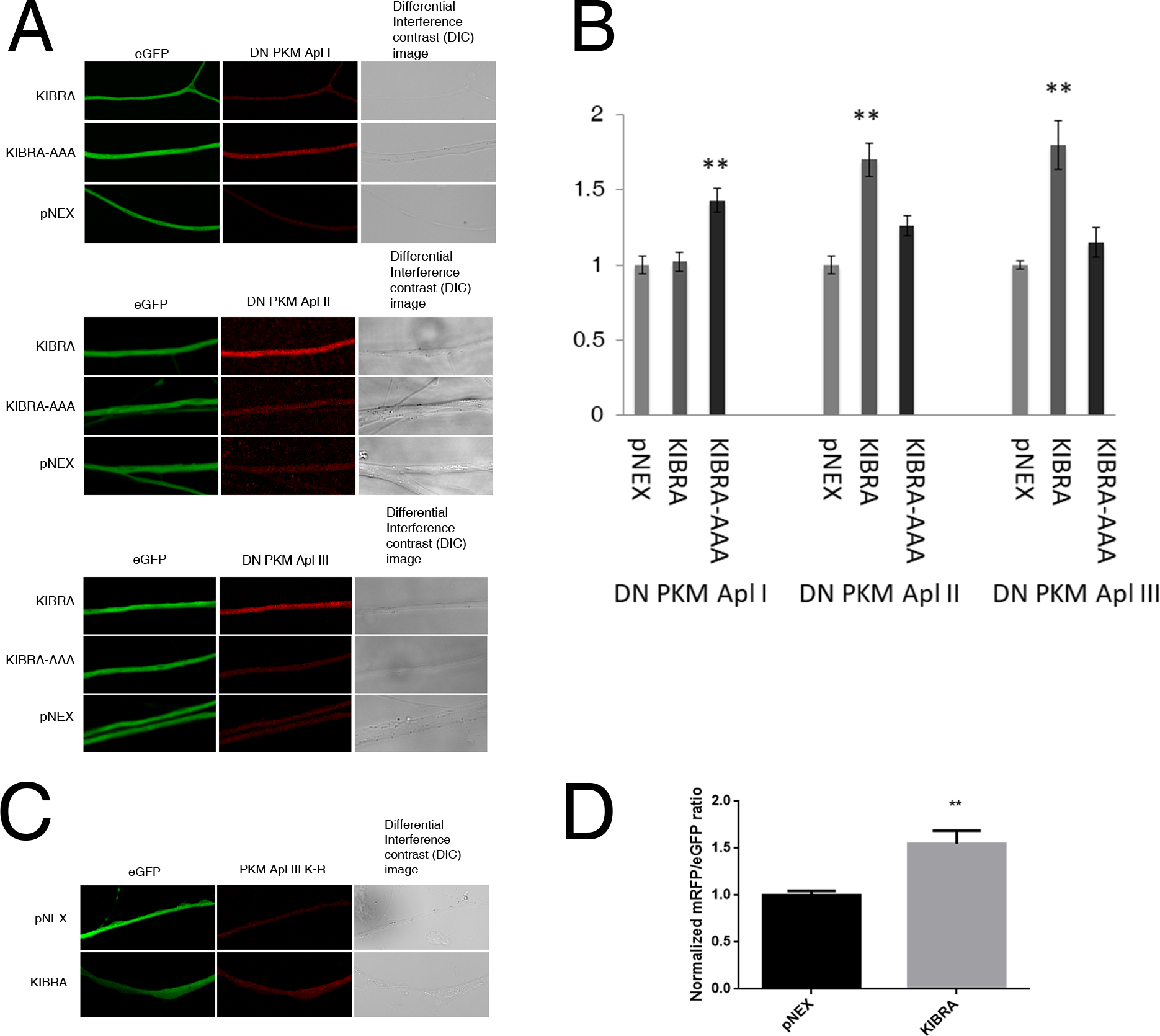
KIBRA stabilizes inactive PKMs in isolated *Aplysia* sensory neurons. **A**, Representative images of eGFP and DN PKM Apl I (top), DN PKM Apl II (middle), and DN PKM Apl III (bottom) with either vector (pNEX3), KIBRA, or KIBRA-AAA in neurites of cultured *Aplysia* sensory neurons 24h after injection. **B**, Stabilization of mRFP-tagged DN PKMs is quantified as the average mRFP/eGFP ratio normalized within each experiment to the average mRFP/eGFP ratio of the pNEX group. KIBRA (AAA) stabilizes DN PKM Apl I, while KIBRA stabilizes DN PKM Apl II and DN PKM Apl III. ANOVAs were performed separately for DN PKM Apl I (pNEX, n=39 neurons; KIBRA, n=66 neurons; KIBRA_RSR-AAA_, n=70 neurons, neurons from four independent experiments; One-way ANOVA with Tukey’s post-Hoc test *), DN PKM Apl II (pNEX, n=61 neurons; KIBRA, n=43 neurons; KIBRA_RSR-AAA_, n=31 neurons, neurons from three independent experiments; One-way ANOVA with Tukey’s post-Hoc test *), and DN PKM Apl III (pNEX, n=28 neurons; KIBRA, n=36 neurons; KIBRA_RSR-AAA_, n=24 neurons, neurons from three independent experiments; One-way ANOVA with Tukey’s post-Hoc test *, p<0.01). **C**, Representative images of eGFP and mRFP-tagged PKM Apl III K-R with either pNEX or KIBRA in neurites of cultured *Aplysia* sensory neurons 24h after injection. **D**, Stabilization of PKM Apl III K-R is quantified as the average mRFP/eGFP ratio normalized as above. PKM Apl III K-R levels are higher in the presence of KIBRA compared to pNEX control (KIBRA, n=31 neurons; pNEX, n=24 neurons, neurons from three independent experiments) * p<0.01, Unpaired Students t-test. Error bars represent SEM.

KIBRA-AAA stabilizes PKM Apl II, but overexpression of KIBRA-AAA in the postsynaptic motor neuron results in a reversal of associative LTF (Hu et al., 2017b) suggesting that PKM Apl II is not involved in associative LTF. However, DN PKM Apl II was able to reverse associative LTF (Hu et al., 2017a). One explanation for this discrepancy is that the overexpressed DN PKM Apl II is competing with endogenous PKM Apl III for KIBRA binding, resulting in a decrease in synaptic strength due to the destabilization of endogenous PKM Apl III. To test this specifically, we overexpressed PKM Apl III along with KIBRA or empty pNEX control and looked at whether concurrent overexpression of DN PKM Apl II affected KIBRA-mediated PKM Apl III stabilization. We also overexpressed DN PKM Apl I to confirm that this competition for KIBRA binding is specific for DN PKM Apl II, as DN PKM Apl I is not stabilized by KIBRA (Fig. 5). As both the DN constructs and the PKM Apl III construct were tagged with mRFP, we quantified PKM Apl III levels using a PKC Apl III C-terminal antibody (Bougie et al, 2009). Overexpression of DN PKM Apl I did not affect KIBRA’s stabilization of PKM Apl III, whereas the stabilization of PKM Apl III in the presence of DN PKM Apl II was significantly reduced and indeed, not significantly different from the vector control (Fig. 6A-B). This demonstrates that DN PKM Apl II is able to interfere with KIBRA’s stabilization of PKM Apl III, supporting the hypothesis that the reversal of associative LTF when DN PKM Apl II is overexpressed in the motor neuron (Hu et al., 2017b) is due to competition with PKM Apl III for KIBRA-mediated stabilization and not the DN’s influence on endogenous PKM Apl II.

**Figure 6.**
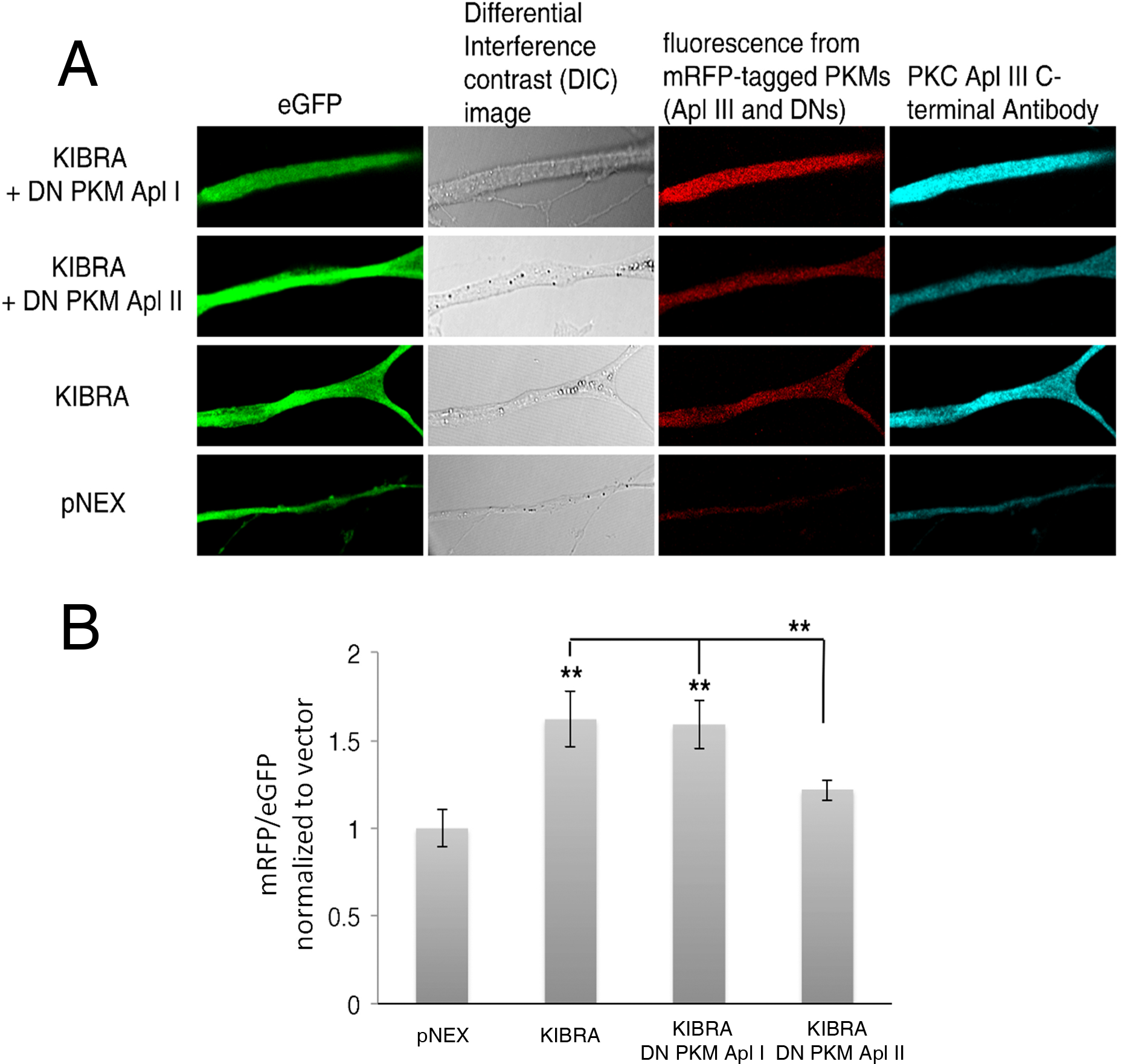
Overexpression of DN PKM Apl II interferes with KIBRA-mediated stabilization of overexpressed PKM Apl III. **A**, Representative images of cultured *Aplysia* sensory neuron neurites co-expressing eGFP, mRFP-tagged PKM Apl III, mRFP-tagged DN PKM Apl I or DN PKM Apl II, and either KIBRA or empty vector (pNEX3). Cultures were fixed and stained with PKC Apl III C-terminal antibody (cyan) prior to imaging. **B**, PKM Apl III stabilization is quantified as the ratio between eGFP fluorescence and PKC Apl III C-terminal antibody staining (cyan) normalized to vector alone. Both KIBRA and KIBRA + DN PKM Apl I stabilized PKM Apl III compared to pNEX control and KIBRA + DN PKM Apl II (One-way ANOVA with Tukey’s post-Hoc test *, p<0.01). [pNEX, n=29 neurons; KIBRA, n=27 neurons; KIBRA + DN PKM Apl I, n=31 neurons; KIBRA + DN PKM Apl II, n=39 neurons, neurons from four independent experiments]. Error bars represent SEM.

### A KIBRA splice variant stabilizes PKM Apl I but not PKM Apl III

The mutant KIBRA (KIBRA-AAA) is able to stabilize PKM Apl I when both are overexpressed in cultured sensory neurons, a stabilization that is not seen with the unmutated KIBRA (Hu et al., 2017b). We hypothesized that this mutation led to a conformational change within the KIBRA protein that opened up a binding site for PKM Apl I. This suggests that the region of KIBRA that binds to PKMs/PKCs is not a peptide, such as a pseudosubstrate, but a domain. Indeed, examination of homology among KIBRAs over evolution reveals three homology regions: the WW domains, the C2 domain, and a third region beginning before the sequence identified as binding to PKMs and spanning approximately 200 amino acids. This region is strongly conserved in all three families of bilaterians (Fig. 7A) and shows around the same percent identity as the WW and C2 domains (Fig. 7B). This region can also be identified in prebilaterians, such as the sea anemone Nematostella, but compared to the C2 and WW domains the new region is poorly conserved in pre-bilaterians (Fig. 7B). Nevertheless, the amino acids required for PKMζ binding are still present in the prebilaterians (Fig. 7A). Interestingly, KIBRA is not present in earlier prebilaterians (sponge and Trichoplax) that lack nervous systems (see Methods). In *Aplysia*, the new region also includes an alternative splicing event that is conserved throughout molluscs (Fig. 7A). This region also contains two alternative inserts in human KIBRA, although not in the same location as the alternative splice insert in *Aplysia* (**purple in** Fig. 7A). To determine if this splicing event could explain isoform specificity of KIBRA binding, we overexpressed a construct containing this splice variant (KIBRA SPL) and assessed the ability of this isoform to stabilize PKM Apl I and PKM Apl III in cultured *Aplysia* sensory neurons. KIBRA SPL does not stabilize PKM Apl III, but it does stabilize PKM Apl I, a pattern of stabilization identical to that of our KIBRA-AAA mutant (Fig.7C). This suggests that production of this splice variant of KIBRA can lead to stabilization of the PKM Apl I isoform, which may be important for the maintenance of non-associative LTF.

**Figure 7.**
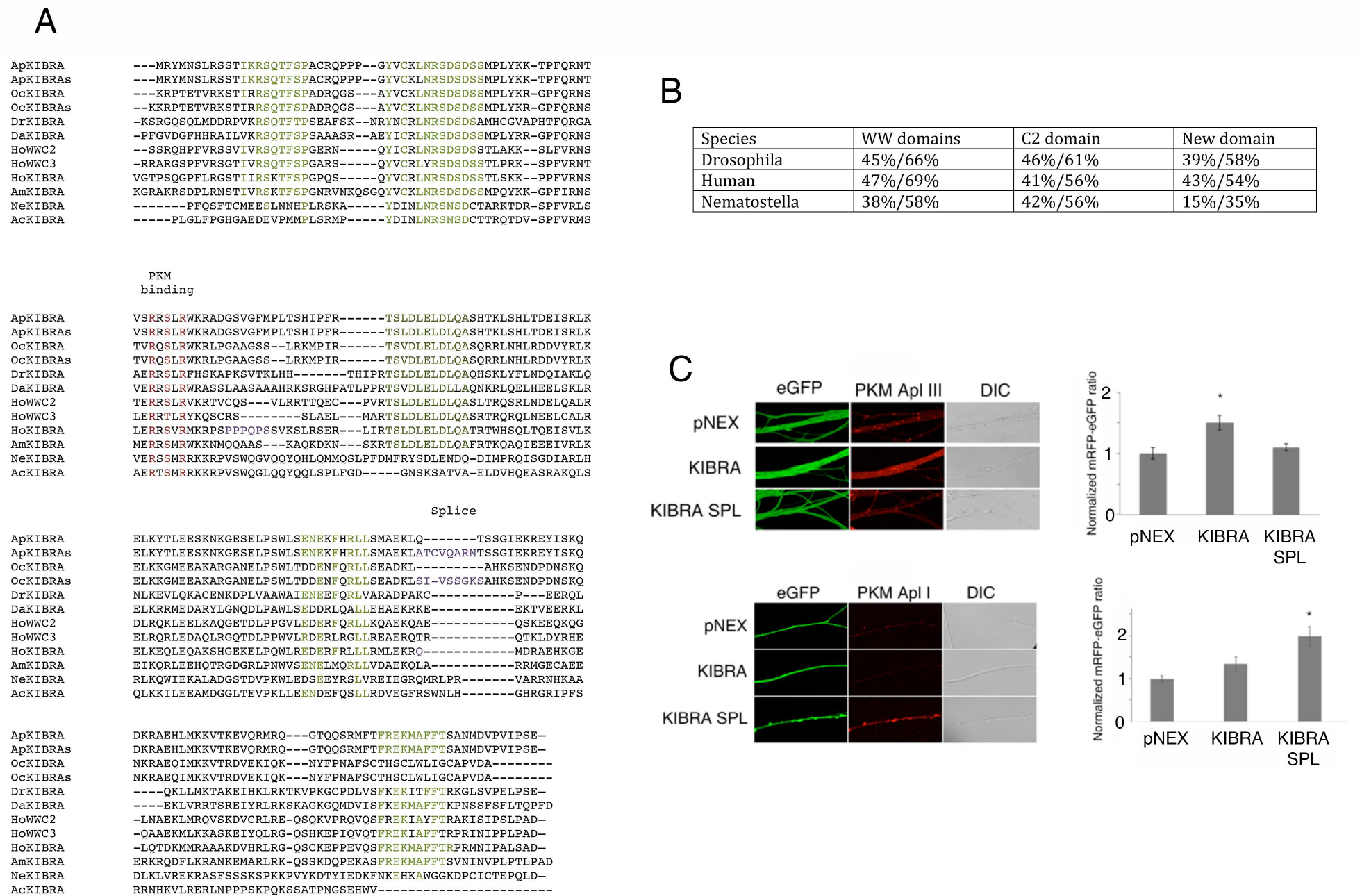
KIBRA Splice (SPL) stabilizes PKM Apl I, but not PKM Apl III. **A**, Alignment of the new domain identified surrounding the PKM binding domain from representative bilaterian and pre-bilaterian animals. Regions highlighted in purple represent alternative mRNA sequences identified by bioinformatics (accession numbers below). The three amino acids required for PKM binding are highlighted in red. Five regions of increased homology are highlighted in green. In vertebrates, the KIBRA gene has two additional paralogues, termed WWC2 and WWC3 (KIBRA’s alternative name is WWC1) and all three are included in the alignment. The sequences were derived from the following sources: Molluscs { Aplysia (Ap) <u>XP_012936697.1</u> 1288:1503; Aplysia splice gene tools: Octopus (Oc) <u>XP_014775126.1</u>1038:1245; Octopus splice <u>KOF84653.1</u>723:904}; Ecdysozoa {Drosophila (Dr) <u>NP_001034055.1</u>966:1156; Daphnia (Da) <u>EFX86200.1</u>876:1090}; Deuterostomes {Amphioxus (Am) <u>XP_019625123.1</u> 995:1209; Human (Ho) WWC2 <u>XP_024309993.1</u>, 913:1116 Human WWC3 <u>NP_056506.2</u>891:1090; Human Kibra<u>XP_016864767.1</u>821:1028l Human Kibra lacking splice PPQPS <u>XP_005265907.1</u>, Human kIbra lacking Q <u>XP_011532791.1</u>; Prebilateria {Nematostella (Anemone, Ne); <u>XP_001629271.1</u>911:1121, Acropora (Coral; Ac) <u>XP_015763151.1</u>950:1136. **B**, Table of homology between the *Aplysia* new region and representative species from the two other bilaterian X and a prebilaterian (Nematostella). First number is percent identity and second is percent similarity from Prot blast at NCBI using *Aplysia* sequence as probe. **C**, Representative images of cultured *Aplysia* sensory neurons expressing eGFP, mRFP-tagged PKM Apl III (top) or PKM Apl I (bottom), and either KIBRA, KIBRA SPL, or empty vector (pNEX3). On left, quantification of stabilization of PKM Apl III (top) and PKM Apl I (bottom). All results are normalized within each experiment to the average mRFP/eGFP ratio of vector alone. KIBRA stabilized PKM Apl III compared to pNEX and KIBRA SPL groups (One-way ANOVA with Tukey’s post-Hoc test *p<0.01[pNEX, n=36 neurons; KIBRA, n=50 neurons; KIBRA SPL, n=32 neurons; neurons from three independent experiments]), while KIBRA SPL stabilized PKM Apl I compared to pNEX and KIBRA groups (One-way ANOVA with Tukey’s post-Hoc test *p<0.001 [pNEX, n=21 neurons; KIBRA, n=25 neurons; KIBRA SPL, n=24 neurons; neurons from three independent experiments]). Error bars represent SEM.

### Isoform specificity is determined by the ‘handle’ helix of PKMs

The isoform specificity of stabilization by KIBRA between PKM Apl I and PKM Apl III must be due to specific sequence differences between the isoforms. We examined the catalytic domains of PKCs for conserved isoform specific differences (Fig. 8A) that were available for binding based on the known crystal structures of these domains (Fig. 8B). We chose the C-terminal (CT) region, already known to bind to proteins with PDZ domains such as PICK 1 (Wan et al., 2012) and an alpha-helical region (helix 5 or helix G in the catalytic domain structure) that appears as a handle in the crystal structures of PKMs (Fig. 8B). We then made chimeras, exchanging these two regions (highlighted in green in Fig. 8A) from the PKM Apl III sequence for the corresponding PKM Apl I sequence, generating PKM Apl III-CT PKM Apl I and PKM Apl III-handle PKM Apl I. We then examined the ability of KIBRA and KIBRA SPL to stabilize these chimeric PKMs. PKM Apl III-CT PKM Apl I behaved like PKM Apl III in that it was stabilized by KIBRA but not by KIBRA SPL (Fig. 8 C-D). In contrast, when the handle region of PKM Apl III was replaced with the handle region of PKM Apl I, stabilization was similar to PKM Apl I—the chimera was stabilized by KIBRA SPL but not by KIBRA (Fig. 8 C-D). While the chimeras were expressed at lower levels than the wild type PKM Apl III (Fig. 8E), the differential stabilization by the two KIBRA splice isoforms indicates that the level of expression does not explain the lack of stabilization. These results indicate that isoform specificity of PKMs is determined by the sequence in the handle helix.

**Figure 8.**
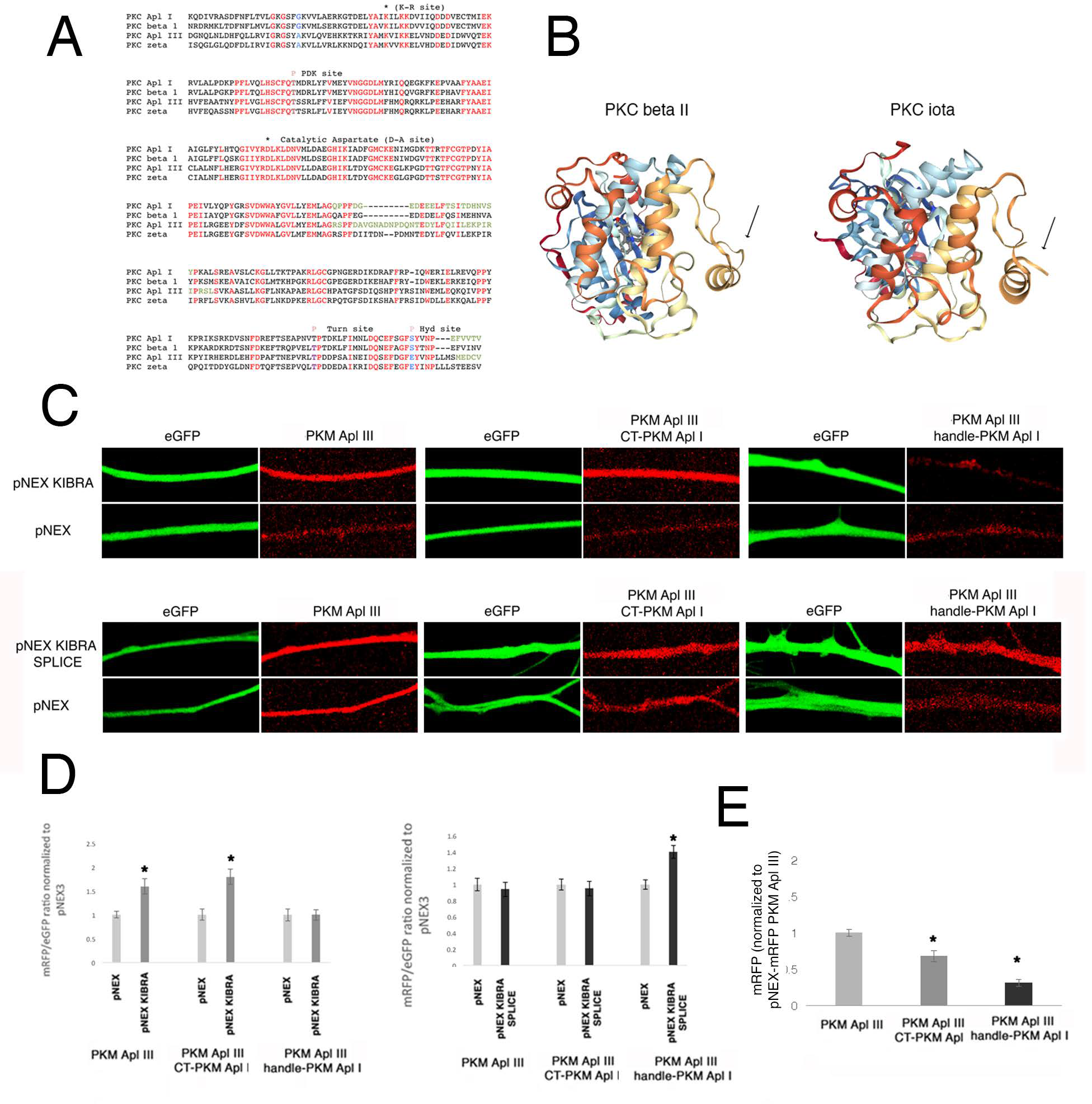
The ‘handle’ domain of PKMs determines isoform specific stabilization by KIBRA. **A**, Sequences of catalytic domains from classical and atypical PKCs from *Aplysia* and human are shown to illustrate regions of homology. Red amino acids are conserved between isoforms. Blue amino acids represent known important differences, a glycine in the ATP binding domain that partially explains the difference in ATP based inhibitors between atypical and classical PKCs and the carboxy-terminal phosphorylation site that is a glutamic acid in atypical PKCs. The sites mutated to form catalytically inactive dominant negatives are illustrated. The green residues are the amino acids removed from PKM Apl III and replaced by the green residues in PKM Apl I in the chimeras. **B**, Rotated structures of the human atypical PKC iota isoform (Messerschmidt et al., 2005) and classical PKC beta II isoform (Grodsky et al., 2006) to illustrate the alpha helix switched in the chimera. From this orientation the helix appears as a ‘handle’. **C**, Representative images of cultured *Aplysia* sensory neurons expressing eGFP, KIBRA or empty vector (pNEX3), and either mRFP-tagged PKM Apl III, PKM Apl III-CT PKM Apl I, or PKM Apl III-handle PKM Apl I. The ability of KIBRA to stabilize the different PKMs was determined by a Student paired T-test between mRFP/eGFP ratios normalized to the pNEX control for each construct. Results were corrected by a Bonferroni test for multiple t-tests (6 for experiments in C and D) *p<0.01. [PKM Apl III, pNEX n=30 neurons, KIBRA n=15 neurons; PKM Apl III-CT PKM Apl I, pNEX n=17 neurons, KIBRA, n=32 neurons; PKM Apl III-handle PKM Apl I, pNEX n=14 neurons, KIBRA n=33 neurons; results are from three independent experiments with all six groups). **D**, Representative images of cultured *Aplysia* sensory neurons expressing eGFP, KIBRA SPL or empty vector (pNEX3), and either PKM Apl III, PKM Apl III-CT PKM Apl I, or PKM Apl III-handle PKM Apl I. The ability of KIBRA SPL to stabilize the different PKMs was determined by a Student paired T-test between mRFP/eGFP ratios normalized to the pNEX control for each construct. Results were corrected by a Bonferroni test for multiple t-tests, p<0.01. [PKM Apl III, pNEX n=23 neurons, KIBRA SPL n=19 neurons; PKM Apl III-CT PKM Apl I, pNEX n=33 neurons, KIBRA SPL, n=23 neurons; PKM Apl III-handle PKM Apl I, pNEX n=20 neurons, KIBRA SPL n=29 neurons; results are from three independent experiments with all six groups). E) The levels of expression of the three constructs in the absence of KIBRA were quantified. The mRFP values are normalized to levels of PKM Apl III in each experiment. One-way ANOVA with Tukey’s post-Hoc test, *p<0.01[PKM Apl III, n=53 neurons;PKM Apl III-CT PKM Apl I, n=51 neurons; PKM Apl III-handle PKM Apl I, n=34 neurons; results are from six independent experiments with all three groups].

## Discussion

### Sufficiency of PKMs for increasing synaptic strength

We investigated the isoform specificity and sufficiency of PKM in regulating synaptic strength by overexpressing distinct isoforms of PKM, presynaptically and postsynaptically. Postsynaptic, but not presynaptic overexpression of PKM was sufficient to increase synaptic strength independent of isoform identity. The lack of sufficiency of PKMs overexpressed in the presynaptic cell is surprising given the ability of activation of PKC in the sensory neuron to increase transmitter release (Ghirardi et al., 1992; Houeland et al., 2007; Manseau et al., 2001; Sugita et al., 1992; Zhou et al., 2014). This includes an important role for cleavage of PKC Apl I to PKM Apl I in the presynaptic cell during activity-dependent ITF (Farah et al., 2017; Sutton et al., 2004). It is possible that the sufficiency of PKM expression in the presynaptic cell to increase synaptic strength is masked by the toxic effects of overexpressing PKM Apl I or PKM Apl II. This toxicity is also seen in the motor neuron, but is delayed compared to the effects in the sensory neuron. Toxicity may be due to a role of PKMs in inducing apoptosis as cleavage of the novel PKC delta by caspases to form a PKM is proapoptotic in many systems (Reyland, 2007). PKM Apl III overexpression is not toxic in either the sensory or motor neuron, despite a similar level of expression as judged by the mRFP level. This allows for the conclusion that PKM Apl III is sufficient in the postsynaptic cell, but not the presynaptic cell, and this result was replicated with two different motor neurons serving as the postsynaptic cell (Fig. 1,2). Despite it not being sufficient to increase synaptic strength in the presynaptic sensory neuron, there is evidence that PKM Apl III is important for non-associative LTF in the sensory neuron. DN PKM Apl III erases non-associative LTF when expressed in the sensory neuron (Hu et al, 2017a) and presynaptic overexpression of a distinct DN form of PKC Apl III blocks non-associative LTF 48 and 72 h after induction (Fiumara et al., 2015). Presynaptic overexpression of PKC Apl III, which leads to production of PKM Apl III (Bougie et al., 2009), was not sufficient to increase synaptic strength, similar to the present study, but did lead to stronger non-associative LTF (Fiumara et al., 2015). In conclusion, PKM Apl III in the sensory neuron is not sufficient to increase synaptic strength, but is required for mediating non-associative LTF. Whether PKM Apl I and PKM Apl II are sufficient to increase synaptic strength in the presynaptic sensory neuron could not be answered due to their toxicity.

### Requirement for PKMs in both presynaptic and postsynaptic cell for associative LTF

While PKM Apl III postsynaptic overexpression is sufficient to increase synaptic strength, this is not sufficient to induce all the features of associative LTF. Associative LTF [which is induced by presynaptic activity in combination with 5HT application in *Aplysia* and is a cellular analogue of classical-conditioning (Buonomano and Byrne, 1990; Hu and Schacher, 2015)] is characterized by a persistent increase in synaptic strength in conjunction with an attenuation of HSD kinetics (Hu and Schacher, 2015); we have shown that this same feature can be achieved by presynaptic overexpression of PKC Apl I in combination with postsynaptic PKM Apl III overexpression. Moreover, presynaptic overexpression of DN PKMs and DN KIBRA-AAA is sufficient to reverse different types of LTF in an isoform specific manner (Hu et al., 2017a). Thus, isoform specificity of PKMs extends to both the presynaptic and postsynaptic side of a synapse.

### Failure of KIBRA overexpression to prolong m-ITF

While the ability of overexpressed PKM to increase synaptic strength suggests that simple stabilization of the PKM produced by m-ITF (Bougie et al., 2012; Villareal et al., 2009) by KIBRA should have led to increased synaptic strength, it is likely that the levels of PKM produced by stimulation are quite different from those produced through overexpression. Simply because large amounts of PKM are sufficient to increase synaptic strength does not necessarily mean that the amounts of PKM produced endogenously are sufficient to increase synaptic strength even when stabilized by KIBRA. Endogenous levels of PKM probably require interactions with other postsynaptic processes generated by LTF that are not induced by m-ITF to increase synaptic strength during LTF.

### KIBRA asserts isoform specificity of PKMs

A weakness of dominant negative constructs is their possible lack of specificity, but this also allows investigators to probe functional aspects of the proteins examined. Overexpression of DN PKM Apl II in the postsynaptic neuron is sufficient to reverse associative LTF (Hu et al., 2017b), suggesting that PKM Apl II may be important in the maintenance of this type of LTF. In the same paper, it was shown that KIBRA-AAA stabilizes PKM Apl II. One might assume, therefore, that overexpression of KIBRA-AAA in the postsynaptic neuron would not have an effect on associative LTF. However, this is not the case. KIBRA-AAA overexpression is sufficient to reverse associative LTF, suggesting that the ability of DN PKM Apl II overexpression to reverse associative LTF is the result of non-specificity of this construct. Here we explain this lack of specificity by demonstrating that DN PKM Apl II interferes with KIBRA-mediated stabilization of PKM Apl III. While our results are consistent with DN PKMs erasing LTF through competition for KIBRA binding, these experiments do not rule out that other interactions could also be important for the actions of the DN PKMs.

### A new conserved region in KIBRA

Examination of KIBRA conservation over evolution demonstrates strong conservation of a 200 amino acid region. This region includes the amino acids previously determined to be required for binding to PKMζ (Vogt-Eisele et al., 2014), and required for the stabilization of PKM Apl III; however, these residues are not required for stabilization of PKM Apl I (Hu et al., 2017b). Along with its role in stabilizing PKMs and regulating AMPA receptor trafficking, KIBRA is also part of the Hippo signalling pathway important for the regulation of cell proliferation in development and in cancer(Genevet et al., 2010; Yu et al., 2010). While KIBRA’s role in this pathway is mainly to act as a scaffold to localize members of this pathway through its WW domains (Baumgartner et al., 2010; Xiao et al., 2011; Zhang et al., 2014), several interactions have been localized to the C-terminal of the protein, including binding to Merlin and dimerization (Baumgartner et al., 2010; Wennmann et al., 2014). These interactions may require this newly identified region.

### How does KIBRA stabilize PKMs?

While the simplest model would be to assume that KIBRA-mediated stabilization is simply due to binding, this may not be the case. KIBRA stabilizes the large tumor suppressor kinase (Lats) as part of its role in the Hippo pathway, but this stabilization can be dissociated from Lats2 binding to KIBRA as binding requires the WW domains, but stabilization does not (Xiao et al., 2011). It is possible that binding of PKMs to KIBRA may instead be involved in localization of the kinase. There is evidence showing an association between KIBRA and AMPA receptors (Makuch et al., 2011), as well as a general enrichment in the postsynaptic density (Johannsen et al., 2008), suggesting the possibility that KIBRA may help localize PKMs to their synaptic targets in addition to their stabilizing effect. It is not clear if KIBRA splice isoforms show differential localization, or whether they have other distinct protein-protein interactions that play a role in isoform specificity.

We have identified features of PKM that are necessary for KIBRA-mediated stabilization. Neither catalytic activity nor priming phosphorylation are required since KIBRA stabilized a DN PKM Apl III made through a K-R mutation that prevents priming phosphorylation (Bougie et al., 2012; Cameron et al., 2009). Switching of one alpha-helix, that in crystal structures resembles a handle, was sufficient to switch the specificity of stabilization by splice forms of KIBRA, suggesting that this is the key region of PKMs that determines isoform specificity. Interestingly, ICAP and zeta-stat, isoform specific PKC inhibitors, are targeted to this region. This region also may play a role in determining isoform-specific substrate specificity of PKCs (Soriano et al., 2016).

## Conclusions

In summary, PKMs have isoform- and synapse-specific roles in the maintenance of LTF. This isoform specificity may be achieved through selective stabilization by KIBRA. The isoform specificity of KIBRA-mediated stabilization of PKMs depends on the handle helix and is independent of catalytic activity of the kinase. These results contribute to our understanding of the mechanisms underlying memory maintenance as well as the synaptic differences among different types of synaptic plasticity that underlie distinct types of memory.

## Acknowledgements

This work is supported by CIHR grant 340328 to WSS, F.S.B Miller Fellowship to LFNational Institutes of Health (NIH) Grant MH-060387 to JH and SS. NIH grants R01 NS02956 and R01 MH096120, and NSF grant IOS 1121690 to DLG.WSS is a James McGill Professor.

## References

Baumgartner, R., I. Poernbacher, N. Buser, E. Hafen, and H. Stocker. 2010. The WW domain protein Kibra acts upstream of Hippo in Drosophila. Dev Cell. 18:309–316.

Bougie, J.K., D. Cai, M. Hastings, C.A. Farah, S. Chen, X. Fan, P.K. McCamphill, D.L. Glanzman, and W.S. Sossin. 2012. Serotonin-induced cleavage of the atypical protein kinase C Apl III in Aplysia. J Neurosci. 32:14630–14640.

Bougie, J.K., T. Lim, C.A. Farah, V. Manjunath, I. Nagakura, G.B. Ferraro, and W.S. Sossin. 2009. The atypical protein kinase C in Aplysia can form a protein kinase M by cleavage. J Neurochem. 109:1129–1143.

Buonomano, D.V., and J.H. Byrne. 1990. Long-term synaptic changes produced by a cellular analog of classical conditioning in Aplysia. Science. 249:420–423.

Cameron, A.J., C. Escribano, A.T. Saurin, B. Kostelecky, and P.J. Parker. 2009. PKC maturation is promoted by nucleotide pocket occupation independently of intrinsic kinase activity. Nat Struct Mol Biol. 16:624–630.

Chen, S., D. Cai, K. Pearce, P.Y. Sun, A.C. Roberts, and D.L. Glanzman. 2014. Reinstatement of long-term memory following erasure of its behavioral and synaptic expression in Aplysia. Elife. 3:e03896.

Dunn, T.W., and W.S. Sossin. 2013. Inhibition of the Aplysia sensory neuron calcium current with dopamine and serotonin. J Neurophysiol. 110:2071–2081.

Farah, C.A., M.H. Hastings, T.W. Dunn, K. Gong, D. Baker-Andresen, and W.S. Sossin. 2017. A PKM generated by calpain cleavage of a classical PKC is required for activity-dependent intermediate-term facilitation in the presynaptic sensory neuron of Aplysia. Learn Mem. 24:1–13.

Fiumara, F., P. Rajasethupathy, I. Antonov, S. Kosmidis, W.S. Sossin, and E.R. Kandel. 2015. MicroRNA-22 Gates Long-Term Heterosynaptic Plasticity in Aplysia through Presynaptic Regulation of CPEB and Downstream Targets. Cell Rep. 11:1866–1875.

Genevet, A., M.C. Wehr, R. Brain, B.J. Thompson, and N. Tapon. 2010. Kibra is a regulator of the Salvador/Warts/Hippo signaling network. Dev Cell. 18:300–308.

Ghirardi, M., O. Braha, B. Hochner, P.G. Montarolo, E.R. Kandel, and N. Dale. 1992. Roles of PKA and PKC in facilitation of evoked and spontaneous transmitter release at depressed and nondepressed synapses in Aplysia sensory neurons. Neuron. 9:479–489.

Greenberg, S.M., V.F. Castellucci, H. Bayley, and J.H. Schwartz. 1987. A molecular mechanism for long-term sensitization in Aplysia. Nature. 329:62–65.

Grodsky, N., Y. Li, D. Bouzida, R. Love, J. Jensen, B. Nodes, J. Nonomiya, and S. Grant. 2006. Structure of the catalytic domain of human protein kinase C beta II complexed with a bisindolylmaleimide inhibitor. Biochemistry. 45:13970–13981.

Hastings, M.H., A. Qiu, C. Zha, C.A. Farah, Y. Mahdid, L. Ferguson, and W.S. Sossin. 2018. The zinc fingers of the small optic lobes calpain bind polyubiquitin. J Neurochem. 146:429–445.

Heitz, F.D., M. Farinelli, S. Mohanna, M. Kahn, K. Duning, M.C. Frey, H. Pavenstadt, and I.M. Mansuy. 2016. The memory gene KIBRA is a bidirectional regulator of synaptic and structural plasticity in the adult brain. Neurobiol Learn Mem. 135:100–114.

Houeland, G., A. Nakhost, W.S. Sossin, and V.F. Castellucci. 2007. PKC modulation of transmitter release by SNAP-25 at sensory-to-motor synapses in aplysia. J Neurophysiol. 97:134–143.

Hu, J., K. Adler, C.A. Farah, M.H. Hastings, W.S. Sossin, and S. Schacher. 2017a. Cell-Specific PKM Isoforms Contribute to the Maintenance of Different Forms of Persistent Long-Term Synaptic Plasticity. J Neurosci. 37:2746–2763.

Hu, J., L. Ferguson, K. Adler, C.A. Farah, M.H. Hastings, W.S. Sossin, and S. Schacher. 2017b. Selective Erasure of Distinct Forms of Long-Term Synaptic Plasticity Underlying Different Forms of Memory in the Same Postsynaptic Neuron. Curr Biol. 27:1888–1899 e1884.

Hu, J., and S. Schacher. 2015. Persistent Associative Plasticity at an Identified Synapse Underlying Classical Conditioning Becomes Labile with Short-Term Homosynaptic Activation. J Neurosci. 35:16159–16170.

Johannsen, S., K. Duning, H. Pavenstadt, J. Kremerskothen, and T.M. Boeckers. 2008. Temporal-spatial expression and novel biochemical properties of the memory-related protein KIBRA. Neuroscience. 155:1165–1173.

Kaang, B.K. 1996. Parameters influencing ectopic gene expression in Aplysia neurons. Neurosci Lett. 221:29–32.

Lisman, J. 1994. The CaM kinase II hypothesis for the storage of synaptic memory. Trends Neurosci. 17:406–412.

Livak, K.J., and T.D. Schmittgen. 2001. Analysis of relative gene expression data using real-time quantitative PCR and the 2(-Delta Delta C(T)) Method. Methods. 25:402–408.

Makuch, L., L. Volk, V. Anggono, R.C. Johnson, Y. Yu, K. Duning, J. Kremerskothen, J. Xia, K. Takamiya, and R.L. Huganir. 2011. Regulation of AMPA receptor function by the human memory-associated gene KIBRA. Neuron. 71:1022–1029.

Manseau, F., X. Fan, T. Hueftlein, W. Sossin, and V.F. Castellucci. 2001. Ca2+-independent protein kinase C Apl II mediates the serotonin-induced facilitation at depressed aplysia sensorimotor synapses. J Neurosci. 21:1247–1256.

McCamphill, P.K., C.A. Farah, M.N. Anadolu, S. Hoque, and W.S. Sossin. 2015. Bidirectional regulation of eEF2 phosphorylation controls synaptic plasticity by decoding neuronal activity patterns. J Neurosci. 35:4403–4417.

Messerschmidt, A., S. Macieira, M. Velarde, M. Badeker, C. Benda, A. Jestel, H. Brandstetter, T. Neuefeind, and M. Blaesse. 2005. Crystal structure of the catalytic domain of human atypical protein kinase C-iota reveals interaction mode of phosphorylation site in turn motif. J Mol Biol. 352:918–931.

Milnik, A., A. Heck, C. Vogler, H.J. Heinze, D.J. de Quervain, and A. Papassotiropoulos. 2012. Association of KIBRA with episodic and working memory: a meta-analysis. Am J Med Genet B Neuropsychiatr Genet. 159B:958–969.

Papassotiropoulos, A., D.A. Stephan, M.J. Huentelman, F.J. Hoerndli, D.W. Craig, J.V. Pearson, K.D. Huynh, F. Brunner, J. Corneveaux, D. Osborne, M.A. Wollmer, A. Aerni, D. Coluccia, J. Hanggi, C.R. Mondadori, A. Buchmann, E.M. Reiman, R.J. Caselli, K. Henke, and D.J. de Quervain. 2006. Common Kibra alleles are associated with human memory performance. Science. 314:475–478.

Reyland, M.E. 2007. Protein kinase Cdelta and apoptosis. Biochem Soc Trans. 35:1001–1004.

Sacktor, T.C. 2011. How does PKMzeta maintain long-term memory? Nat Rev Neurosci. 12:9–15.

Shema, R., S. Haramati, S. Ron, S. Hazvi, A. Chen, T.C. Sacktor, and Y. Dudai. 2011. Enhancement of consolidated long-term memory by overexpression of protein kinase Mzeta in the neocortex. Science. 331:1207–1210.

Soriano, E.V., M.E. Ivanova, G. Fletcher, P. Riou, P.P. Knowles, K. Barnouin, A. Purkiss, B. Kostelecky, P. Saiu, M. Linch, A. Elbediwy, S. Kjaer, N. O’Reilly, A.P. Snijders, P.J. Parker, B.J. Thompson, and N.Q. McDonald. 2016. aPKC Inhibition by Par3 CR3 Flanking Regions Controls Substrate Access and Underpins Apical-Junctional Polarization. Dev Cell. 38:384–398.

Sossin, W.S. 2007. Isoform specificity of protein kinase Cs in synaptic plasticity. Learn Mem. 14:236–246.

Sossin, W.S. 2018. Memory Synapses Are Defined by Distinct Molecular Complexes: A Proposal. Front Synaptic Neurosci. 10:5.

Sugita, S., J.R. Goldsmith, D.A. Baxter, and J.H. Byrne. 1992. Involvement of protein kinase C in serotonin-induced spike broadening and synaptic facilitation in sensorimotor connections of Aplysia. J Neurophysiol. 68:643–651.

Sutton, M.A., M.W. Bagnall, S.K. Sharma, J. Shobe, and T.J. Carew. 2004. Intermediate-term memory for site-specific sensitization in aplysia is maintained by persistent activation of protein kinase C. J Neurosci. 24:3600–3609.

Sutton, M.A., J. Ide, S.E. Masters, and T.J. Carew. 2002. Interaction between amount and pattern of training in the induction of intermediate- and long-term memory for sensitization in aplysia. Learn Mem. 9:29–40.

Tsokas, P., C. Hsieh, Y. Yao, E. Lesburgueres, E.J.C. Wallace, A. Tcherepanov, D. Jothianandan, B.R. Hartley, L. Pan, B. Rivard, R.V. Farese, M.P. Sajan, P.J. Bergold, A.I. Hernandez, J.E. Cottrell, H.Z. Shouval, A.A. Fenton, and T.C. Sacktor. 2016. Compensation for PKMzeta in long-term potentiation and spatial long-term memory in mutant mice. Elife. 5.

Villareal, G., Q. Li, D. Cai, A.E. Fink, T. Lim, J.K. Bougie, W.S. Sossin, and D.L. Glanzman. 2009. Role of protein kinase C in the induction and maintenance of serotonin-dependent enhancement of the glutamate response in isolated siphon motor neurons of Aplysia californica. J Neurosci. 29:5100–5107.

Vogt-Eisele, A., C. Kruger, K. Duning, D. Weber, R. Spoelgen, C. Pitzer, C. Plaas, G. Eisenhardt, A. Meyer, G. Vogt, M. Krieger, E. Handwerker, D.O. Wennmann, T. Weide, B.V. Skryabin, M. Klugmann, H. Pavenstadt, M.J. Huentelmann, J. Kremerskothen, and A. Schneider. 2014. KIBRA (KIdney/BRAin protein) regulates learning and memory and stabilizes Protein kinase Mzeta. J Neurochem. 128:686–700.

Wan, Q., X.Y. Jiang, A.M. Negroiu, S.G. Lu, K.S. McKay, and T.W. Abrams. 2012. Protein kinase C acts as a molecular detector of firing patterns to mediate sensory gating in Aplysia. Nat Neurosci. 15:1144–1152.

Wennmann, D.O., J. Schmitz, M.C. Wehr, M.P. Krahn, N. Koschmal, S. Gromnitza, U. Schulze, T. Weide, A. Chekuri, B.V. Skryabin, V. Gerke, H. Pavenstadt, K. Duning, and J. Kremerskothen. 2014. Evolutionary and molecular facts link the WWC protein family to Hippo signaling. Mol Biol Evol. 31:1710–1723.

Xiao, L., Y. Chen, M. Ji, and J. Dong. 2011. KIBRA regulates Hippo signaling activity via interactions with large tumor suppressor kinases. J Biol Chem. 286:7788–7796.

Yao, Y., M.T. Kelly, S. Sajikumar, P. Serrano, D. Tian, P.J. Bergold, J.U. Frey, and T.C. Sacktor. 2008. PKM zeta maintains late long-term potentiation by N-ethylmaleimide-sensitive factor/GluR2-dependent trafficking of postsynaptic AMPA receptors. J Neurosci. 28:7820–7827.

Yoshihama, Y., T. Hirai, T. Ohtsuka, and K. Chida. 2009. KIBRA Co-localizes with protein kinase Mzeta (PKMzeta) in the mouse hippocampus. Biosci Biotechnol Biochem. 73:147–151.

Yu, J., Y. Zheng, J. Dong, S. Klusza, W.M. Deng, and D. Pan. 2010. Kibra functions as a tumor suppressor protein that regulates Hippo signaling in conjunction with Merlin and Expanded. Dev Cell. 18:288–299.

Zhang, L., S. Yang, D.O. Wennmann, Y. Chen, J. Kremerskothen, and J. Dong. 2014. KIBRA: In the brain and beyond. Cell Signal. 26:1392–1399.

Zhao, Y., K. Leal, C. Abi-Farah, K.C. Martin, W.S. Sossin, and M. Klein. 2006. Isoform specificity of PKC translocation in living Aplysia sensory neurons and a role for Ca2+-dependent PKC APL I in the induction of intermediate-term facilitation. J Neurosci. 26:8847–8856.

Zhou, L., D.A. Baxter, and J.H. Byrne. 2014. Contribution of PKC to the maintenance of 5-HT-induced short-term facilitation at sensorimotor synapses of Aplysia. J Neurophysiol. 112:1936–1949.

